# CRISPR enriches for cells with mutations in a p53-related interactome, and this can be inhibited

**DOI:** 10.1101/2021.03.10.434760

**Authors:** Long Jiang, Katrine Ingelshed, Yunbing Shen, Sanjaykumar V. Boddul, Vaishnavi Srinivasan Iyer, Zsolt Kasza, Saikiran Sedimbi, David P. Lane, Fredrik Wermeling

## Abstract

CRISPR/Cas9 can be used to inactivate or modify genes by inducing double-stranded DNA breaks^1–3^. As a protective cellular response, DNA breaks result in p53-mediated cell cycle arrest and activation of cell death programs^4,5^. Inactivating *p53* mutations are the most commonly found genetic alterations in cancer, highlighting the important role of the gene^6–8^. Here, we show that cells deficient in p53, as well as in genes of a core CRISPR-p53 tumor suppressor interactome, are enriched in a cell population when CRISPR is applied. Such enrichment could pose a challenge for clinical CRISPR use. Importantly, we identify that transient p53 inhibition suppresses the enrichment of cells with these mutations. Furthermore, in a data set of >800 human cancer cell lines, we identify parameters influencing the enrichment of *p53* mutated cells, including strong baseline *CDKN1A* expression as a predictor for an active CRISPR-p53 axis. Taken together, our data identify strategies enabling safe CRISPR use.

---

The CRISPR molecular biology tool has immense potential for clinical gene therapy use^9^. Correcting disease-causing mutations in congenital monogenic disorders affecting for example the hematopoietic system are apparent candidates, and clinical trials for sickle cell anemia and beta-thalassemia are ongoing^10–12^. CRISPR-mediated modifications of cells used for chimeric antigen receptor (CAR) based immunotherapy is another clinical setting where CRISPR is likely to have a large impact^13,14^. Concerns about the safety of CRISPR-based gene therapy are successfully being addressed at multiple levels. This includes risks related to CRISPR off-target activity, where, for example, sgRNA design with low off-target activity^15^, high fidelity Cas9 versions^16^, and methods for the evaluation of off-target mutations^17^ has been an intense research focus. Another risk that has been suggested is that the CRISPR-mediated DNA damage could give cells with mutations in *p53* (also referred to as *TP53* in humans, and *Trp53* in mice) a selective advantage and thereby be enriched in a cell population exposed to CRISPR^18–20^. Notably, *TP53* mutations are seen in human embryonic stem cell lines^21^ and also contribute to clonal hematopoiesis^22,23^ showing that premalignant *TP53* mutations can be found in different cell populations of relevance for clinical CRISPR use.

To address the risk of enriching for cells with *p53* mutations using CRISPR, we initially studied the cellular response following CRISPR (electroporation of a GFP targeting sgRNA with very low off-target activity, as calculated by Cas-OFFinder^24^), and compared it to the response following a pulse of Etoposide (a topoisomerase II inhibitor causing DNA damage and p53 activaton^25^), or AMG232 (an inhibitor activating p53 by interfering with the MDM2-p53 interaction^26^). Using a cell line derived from mouse hematopoietic stem cells of Cas9+ GFP+ mice (**Supplementary Fig. 1a-d**), we observed that the CRISPR event resulted in partial delayed cell growth (**Fig. 1b**), apoptosis induction (**Fig. 1c**), and transcription of *Cdkn1a* (also referred to as *p21*, a p53 target gene central to DNA damage-induced cell cycle arrest response^27^, **Fig. 1d**), although at a lower magnitude compared to AMG232 (Fig. 1b), and Etoposide (Fig. 1b-d). To explore if this relatively mild phenotype was sufficient to give a selective advantage to cells with mutations in *Trp53*, we established a competitive assay, where *Trp53* KO and WT cells were mixed at a 1:4 ratio, and subsequently exposed to CRISPR (electroporation or lentiviral delivery of sgRNA), or to pharmacological p53 activation with AMG232 or Etoposide (**Fig. 1e**). Notably, the proportion of *Trp53* KO cells in the population did not expand significantly by only being cultured, or by being transduced by non-targeting control (NTC) lentiviral particles, as identified by sequencing the *Trp53* locus after seven days in culture. However, the proportion of *Trp53* KO cells did significantly expand after being exposed to CRISPR, AMG232, and Etoposide (**Fig. 1f** **and** **Supplementary Fig. 1e**), as well as hypoxia (**Supplementary Fig. 1f-h**). The level of enrichment mirrored the severity of the cellular phenotype shown in figure 1b-d, with Etoposide and AMG232 causing a significantly higher enrichment of *Trp53* KO cells compared to CRISPR. Based on this, we hypothesized that the level of DNA damage response induced by sgRNAs could be a parameter affecting the enrichment of *Trp53* KO cells. To this end, we designed a set of sgRNAs with different off-target activity and used them alone or in combination at equimolar concentrations, and compared the induction of *Cdkn1a* transcription at early time points (0-24h), as a proxy for p53 activation, to the enrichment of cells with *Trp53* mutations at a later time points (seven days). We found that the level of early CRISPR-induced *Cdkn1a* transcription correlated with the enrichment of *Trp53* KO cells, and that the level of enrichment could be estimated based on the level of off-target activity one, or a combination, of sgRNAs would give (**Fig. 1g-h** **and** **Supplementary Fig. 2a-c**). This suggests that the level of induced *Cdkn1a* expression could be a relevant parameter for sgRNA selection.

**Figure 1.**
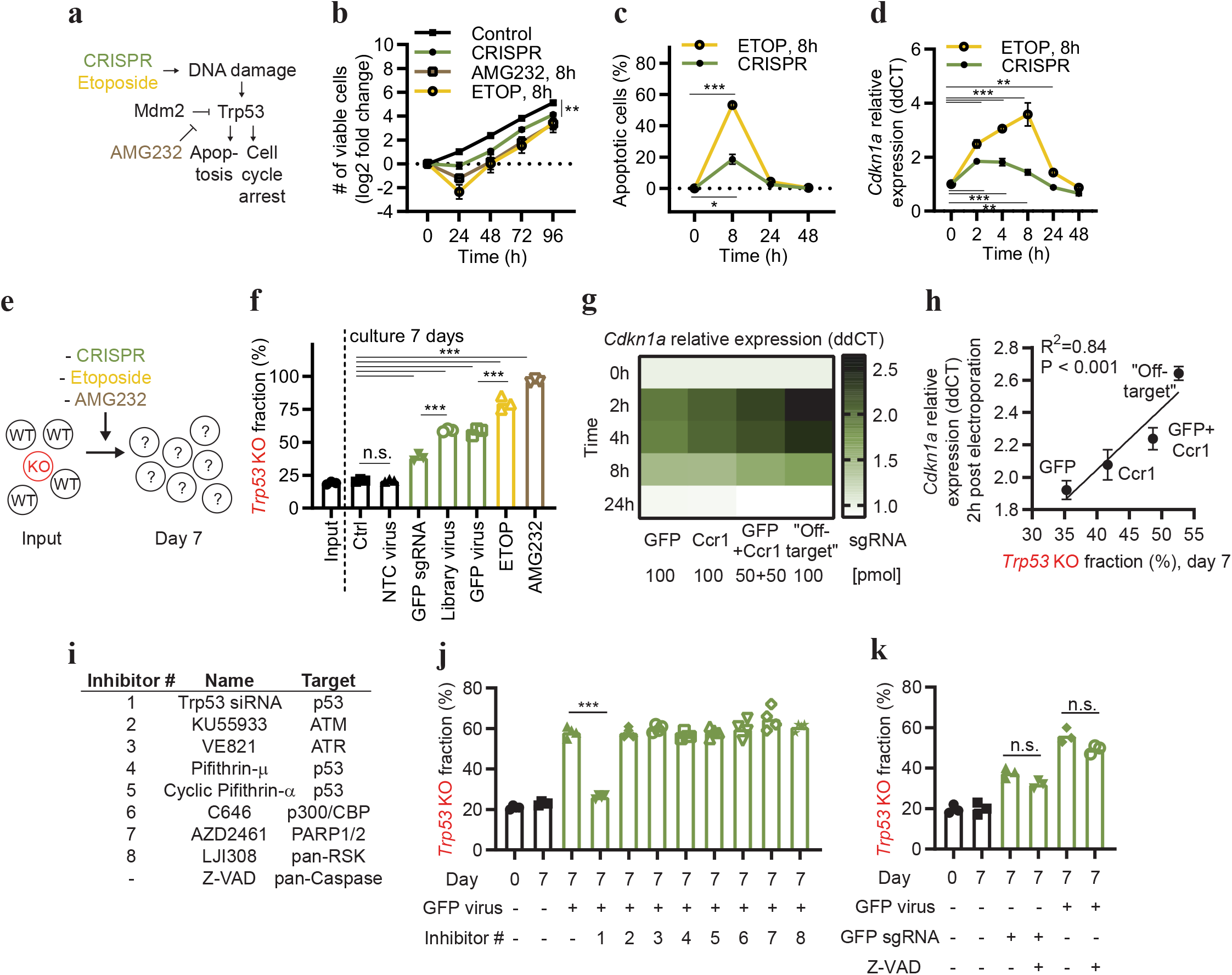
CRISPR-mediated DNA damage enriches for cells with mutations in Trp53, and this can be inhibited. (**a**) Model describing how CRISPR and the used drugs are expected to affect targeted cells. (**b**) Growth characteristics of Hox expanded bone marrow cells (from Cas9+ GFP+ mice), exposed to CRISPR (electroporated with a GFP sgRNA, or control), or pulsed for 8h with AMG232, or Etoposide (ETOP). (**c**) Kinetic analysis of apoptosis by flow cytometry-based TUNEL assay of Hox cells exposed to CRISPR (electroporated with GFP sgRNA) or ETOP. (**d**) Kinetic qPCR analysis of *Cdkn1a* expression of Hox cells exposed to CRISPR (electroporated with GFP sgRNA) or ETOP. (**e**) Model describing experimental setup. (**f**) WT and Trp53 KO Hox cells (Cas9+ and GFP+) were mixed and subjected to CRISPR (electroporated with a GFP sgRNA, non-targeting ctrl (NTC) virus, CRISPR library virus, or GFP targeting sgRNA virus), or an 8h pulse with ETOP or AMG232. After seven days in culture, cells were sequenced, and the fraction of Trp53 KO sequences determined. (**g**) Hox cells (Cas9+ and GFP+) were electroporated with indicated sgRNAs, and *Cdkn1a* expression was analyzed by qPCR at different time points as indicated in figure. (**h**) Comparison of the *Cdkn1a* expression by qPCR 2h post electroporation with indicated sgRNAs, and enrichment of Trp53 KO sequences day seven. (**i**) Inhibitors used in (j-k). (**j**) WT and Trp53 KO Hox cells (Cas9+ and GFP+) were mixed and transduced with a GFP targeting sgRNA virus in the presence of inhibitors. Cells were then cultured for seven days, followed by sequencing of Trp53, and the frequency of Trp53 mutations quantified. (**k**) WT and Trp53 KO Hox cells (Cas9+ and GFP+), were mixed and transduced with a GFP targeting sgRNA virus or electroporated with a GFP sgRNA in the presence of the caspase inhibitor Z-VAD. Cells were then cultured for seven days, followed by sequencing of Trp53, and the frequency of Trp53 mutations quantified. Data is shown as mean +/− SEM, n=3 (b-d, h), mean and individual values, n=3-4 (f, j-k), or heatmap based on the average signal, n=3 (g). Data is representative of two or more experiments. n.s. = non-significant, * = p < 0.05, ** = p < 0.01, *** = p < 0.001 by two-way ANOVA and Turkey’s post-test (b-d), one-way ANOVA and Tukey’s post-test (f, j-k), or Pearson r correlation and simple linear regression line (h).

The enrichment of cells with inactive p53 could pose a challenge to the clinical use of CRISPR. Therefore, we set out to identify strategies that could be used to suppress the enrichment. We hypothesized that inhibiting p53, or proteins playing a non-redundant role in the p53 pathway could be a viable strategy. We gathered a selection of potentially relevant inhibitors (**Fig. 1i**), and pretreated cells with the inhibitors followed by exposing them to CRISPR. Importantly, we found that treating the cells with a Trp53 siRNA completely inhibited the enrichment of cells with *Trp53* mutations, while the other used inhibitors did not show sufficient activity (**Fig. 1j** **and** **Supplementary Fig. 3**). We also noted that the pan-Caspase inhibitor Z-VAD did not significantly suppress the enrichment of cells with *Trp53* mutations (**Fig. 1k**), suggesting that apoptosis is not a significant driver in the CRISPR-mediated enrichment of *Trp53* KO cells. We concluded (*i*) that cells with inactivating *Trp53* mutations are enriched following exposure to CRISPR, (*ii*) that this correlates to the level of triggered DNA damage response, and (*iii*) that transient inhibition of p53 can completely abrogate this enrichment.

Next, we set out to identify p53 linked genes playing a non-redundant role in the CRISPR-induced DNA damage response. Such genes could represent additional drug targets to modify the CRISPR-p53 response and, importantly, cells with mutations in such genes could be enriched by CRISPR-mediated DNA damage. To this end, we applied a custom CRISPR screen library targeting 395 DNA damage-related genes and controls, with 1640 sgRNAs (**Supplementary Table 1**), to WT and *Trp53* KO cells. Relying on the DNA damage response induced by the introduction of the sgRNA library into Cas9+ cells, as has been used in the past^19,28^, we found that Trp53 sgRNAs were enriched and that Mdm2 sgRNAs were depleted in a p53 dependent manner, in both Hox and B16 cells (**Supplementary Fig. 4a-d**). In this regard, *Mdm2* behaved as an essential gene, where mutated cells are rapidly lost over time (**Supplementary Fig. 4e-f**). We also noted that the Hox cells, both *Trp53* WT and KO, but not B16 cells, over time enriched for sgRNAs relating to type I IFN signaling (Stat1, Jak1, and Ptprc), and were sensitive to induced type I IFN signaling (**Supplementary Fig. 4g-j**, **and Supplementary Table 2**).

Next, we performed a screen using the same sgRNA library, but this time culturing the cells for 14 days after the introduction of the library into the Hox cell population, and then applied a controlled CRISPR DNA damage event by electroporating a GFP targeting sgRNA, comparing it to an eight-hour pulse of AMG232 or Etoposide (**Fig. 2a**). In this way, we could separate the studied CRISPR event from the introduction of the CRISPR library. Seven days later we collected the different cell populations, sequenced the sgRNA barcodes, and deconvoluted the sgRNA enrichment/depletion using MAGeCK^29^. Comparing the different interventions to control-treated cells, we identified that CRISPR enriched for cells with mutations in *Chek2*, *Trp53*, and *Cdkn1a* (**Fig. 2b**); AMG232 enriched for cells with mutations in *Trp53* and *Cdkn1a* (**Fig. 2c**); and Etoposide enriched for cells with mutations in *Atm*, *Chek2*, *Trp53* and *Cdkn1a* (**Fig. 2d**). Notably, AMG232 also enriched for mutations in *Stat1*, and *Eif2ak2*, two genes related to type I IFN signaling (Fig. 2c), again highlighting this pathway in the Hox cells. Focusing on the Atm-Chek2-Trp53-Cdkn1a pathway, linking double-stranded DNA damage to cell cycle arrest^4,5,27^, we found that CRISPR caused a significant enrichment of sgRNAs targeting all these genes when comparing WT and *Trp53* KO cells (**Fig. 2e-h**) and that a Trp53 siRNA could significantly suppress the enrichment of cells with mutations in *Chek2*, *Trp53*, and *Cdkn1a*, but not for *Atm* for which the enrichment phenotype also was the weakest (**Fig. 2i** **and** **Supplementary Fig. 5a**). Noteworthy, we did not observe any enrichment of sgRNAs targeting genes related to apoptosis (**Fig. 2j** **and** **Supplementary Fig. 5b**), in line with data presented by Haapaniemi et. al^19^, and the absence of activity of the pan-Caspase inhibitor Z-VAD (Fig. 1k). We concluded (*i*) that the Atm-Chek2-Trp53-Cdkn1a pathway plays a non-redundant role in the CRISPR-mediated DNA damage response (**Fig. 2k**), and as a consequence (*ii*) that cells with mutations in these tumor suppressor genes could be enriched in a cell population as CRISPR is applied, also if they are found at a low frequency (1:1640 in figure 2 compared to 1:4 in figure 1e-j), and (*iii*) that a Trp53 siRNA can suppress the enrichment of mutated cells to a large degree.

**Figure 2.**
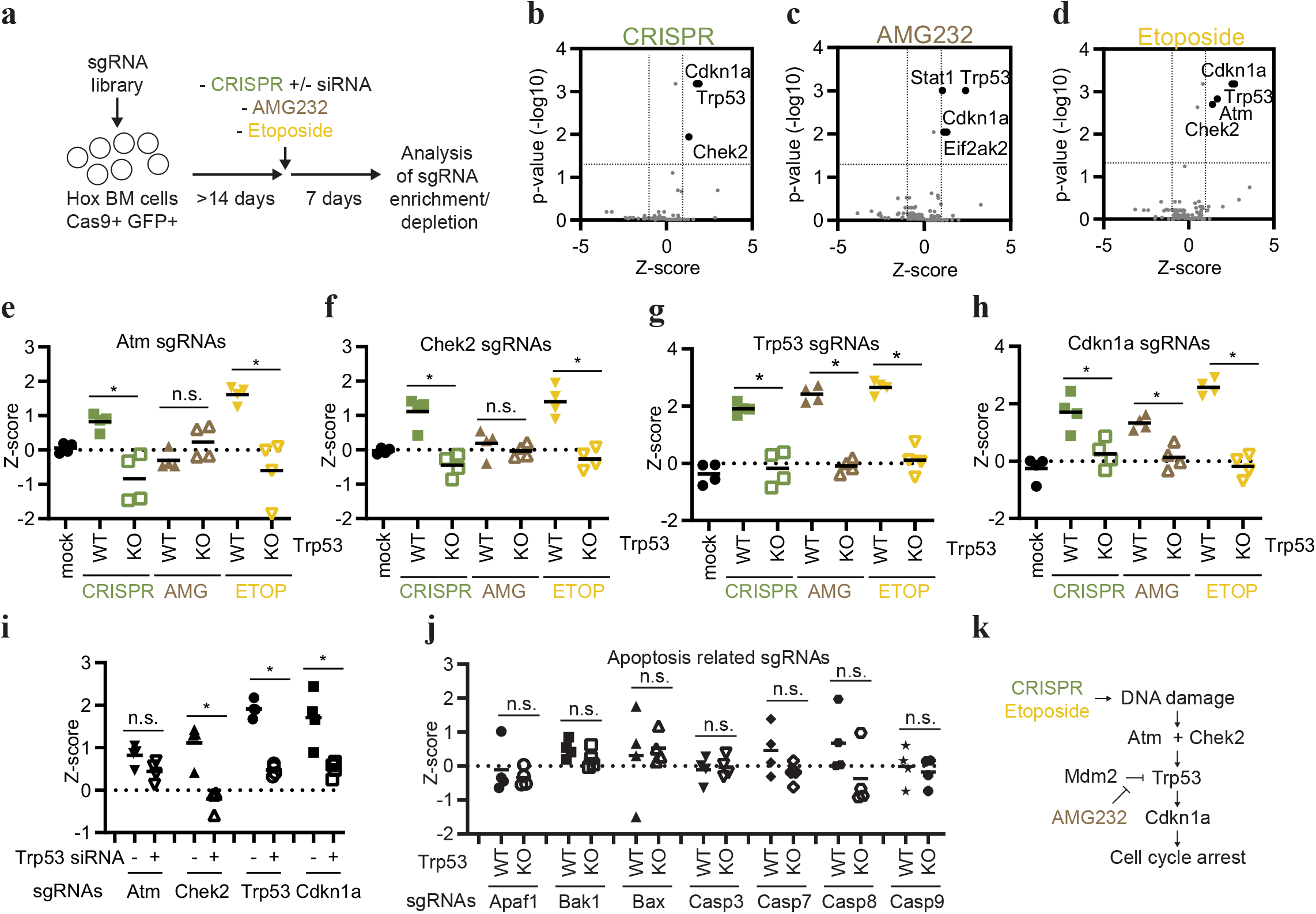
CRISPR enriches for low-frequency mutations in tumor suppressor genes. (**a**) Model describing experimental setup. (**b-d**) Hox cells (Cas9+ and GFP+) were transduced with a custom CRISPR library and cultured for >14 days. Cells were then either exposed to CRISPR (GFP targeting sgRNA) (b), AMG232 8h pulse (c), or Etoposide 8h pulse (d). Cells were cultured for seven days and then the sgRNA representation was analyzed by next-generation sequencing, and enrichment/depletion deconvoluted by MAGeCK. (**e-h**) Z-scores of individual sgRNAs (n=4/gene) for Atm (e), Chek2 (f), Trp53 (g), and Cdkn1a (h) in WT and Trp53 KO Hox cells treated with mock (electroporation without sgRNA), CRISPR (electroporation with GFP targeting sgRNA), AMG232 or ETOP. (**i**) Z-scores of individual sgRNAs (n=4/gene) in Hox cells treated with Trp53 siRNA at the same time as being electroporated with a GFP targeting sgRNA as described in (a). (**j**) Z-scores of individual sgRNAs (n=4/gene) for genes linked to apoptosis. (**k**) Model indicating genes playing a non-redundant role in the DNA damage response. Data presented as volcano plots with Z-score (log2 fold change) and adjusted p-values (b-d), or mean and individual values for 4 gRNAs from the exploratory screen (e-j). * = P < 0.05, and n.s. = non-significant by Mann-Whitney test.

To expand our understanding of which mutations could be enriched in a cell population as CRISPR is applied, we next turned to the Depmap portal, containing full genome CRISPR screen data of 808 human cell lines (Public 20Q4 release), as well as, for example, baseline gene expression, mutation status, and drug sensitivity data for a large proportion of the same cells^30–32^. Exploring the database, we found that 103 of the 808 included cell lines (12.7%) enriched for TP53 sgRNAs as defined by a score >1. Stratifying these cells based on *TP53* mutation status (WT or any type of mutation, making up 32.3% and 67.7% of the cell lines, respectively), we found that 94 of the 103 cell lines (91.3%) that enriched for TP53 sgRNAs were confined to the *TP53* WT group (**Fig. 3a**), supporting the validity of the analysis. As an anecdote, we also noted that the cell line with the strongest enrichment for TP53 sgRNAs is a version of the RPE-1 cell line (**Supplementary Table 3**), that has been used in a significant portion of previous publications related to p53 and CRISPR^19,28,33,34^. Next, we compared the TP53 sgRNA enrichment to the sensitivity of the cells to p53 modulating drugs. We found a clear correlation between TP53 sgRNA enrichment and the sensitivity to AMG232 (**Fig. 3b-c**), as well as to Nutlin-3 and CMG097 (both with a similar mode of action as AMG232), but not, for comparison, to the p53 inhibitor Pifithrin-μ (**Supplementary Fig. 6**). Taken together, we concluded that TP53 sgRNA enrichment in the Depmap dataset could be used to identify cells where the CRISPR-p53 pathway is active, and, as a consequence, that correlation to TP53 sgRNA enrichment also could identify other factors influencing the pathway. In line with this concept, we found that TP53 sgRNA enrichment correlated strongly with MDM2 sgRNA depletion (**Fig. 3d**). Performing the same correlation analysis, but on a full genome basis, we identified a list of genes where sgRNA enrichment (+) or depletion (−) correlated with TP53 sgRNA enrichment (**Fig. 3e** **and Supplementary Table 4**). The list contained the genes we had identified in our experimental data and expanded the list of genes playing a non-redundant, TP53-related role in the CRISPR-mediated DNA damage response. Furthermore, analyzing the top 10 enriched and depleted genes using geneMANIA, we noted a high level of physical interaction between the linked proteins (**Fig. 3f**). An alternative analysis approach of the data, identifying genes where sgRNA enrichment/depletion correlated with the *TP53* mutation status, instead of TP53 sgRNA enrichment (as in Fig. 3e), resulted in a list of genes with a large level of overlap with the list in figure 3e (**Supplementary Fig. 7** **and Supplementary Table 5**). Based on the overlap of these lists, and the experimental data we highlight TP53, TP53BP1, CHEK2, ATM, CDKN1A, USP28, UBE2K, XPO7 as a core CRISPR-p53 tumor suppressor interactome, where cells with inactivating mutations or silencing of these genes (something that is commonly found in cancers^6–8,35–39^, **Supplementary Fig. 8**) can be expected to be enriched in cell population as CRISPR is applied. Furthermore, cells with copy number amplifications, overexpression, or activating mutations of *MDM2*, *PPM1D*, *MDM4*, *DDX31*, *USP7*, *PPM1G*, *WDR89*, and *TERF1* also observed in cancer ^40–45^, could similarly be enriched by CRISPR.

**Figure 3.**
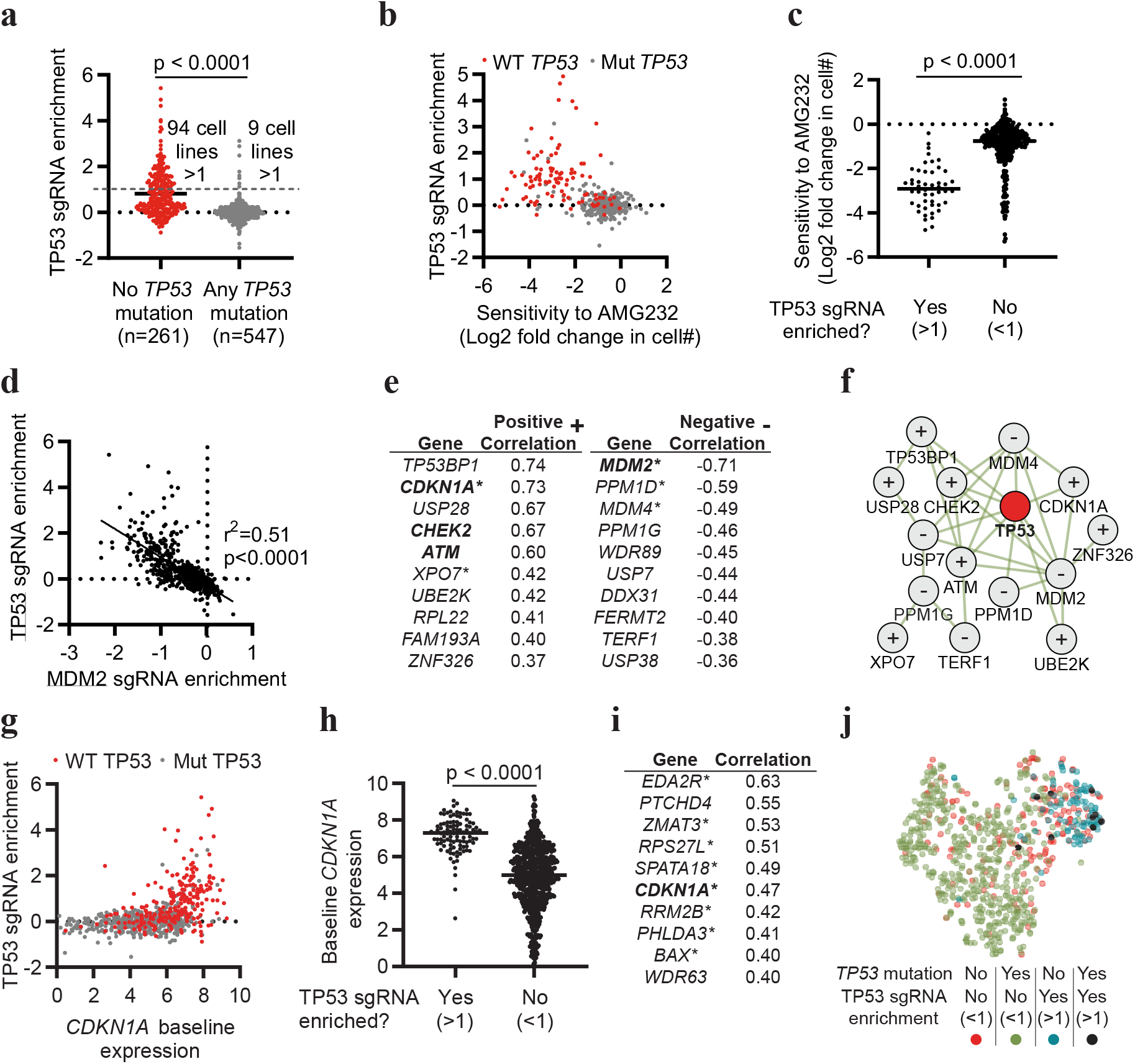
Enrichment of TP53 sgRNAs in full genome CRISPR screens of human cancer cell lines. (**a**) Enrichment score of TP53 gRNAs in 808 cell lines, stratified based on the presence or absence of any mutation in TP53. (**b**) Correlation between the enrichment of TP53 sgRNAs and AMG232 sensitivity. TP53 WT cells are indicated in red. (**c**) Sensitivity to AMG232 in cell lines stratified based on TP53 sgRNA enrichment. (**d**) Correlation between the enrichment of TP53 sgRNAs and MDM2 sgRNAs. (**e**) Top 10 genes with the strongest positive (+) and negative (−) correlation with TP53 sgRNA enrichment from full genome CRISPR screens of 808 cell lines. * indicates genes identified as transcription factor target genes for p53 identified by Enricher. Bold indicates genes identified experimentally in figure 2. (**f**) Physical interactions of genes in (e) defined by geneMANIA. + indicates genes that positively correlate, and - indicates genes that negatively correlate with TP53 sgRNA enrichment. (**g**) Correlation of TP53 sgRNA enrichment to baseline CDKN1A expression. TP53 WT cells are indicated in red. (**h**) Baseline CDKN1A expression in cells stratified based on the enrichment of TP53 sgRNAs. (**i**) Top 10 genes with expression correlating with enrichment of TP53 sgRNAs. * indicates genes identified as transcription factor target genes for p53. (**j**) tSNE dimensionality reduction analysis of cell, n=808, based on expression of the ten genes in (i). Data includes all available data in the Depmap CRISPR (Avana) 20Q4, Expression Public 20Q4, as well as Drug sensitivity (PRISM Repurposing Primary Screen) 19Q4 releases. Each dot represents one cell line (a-d, g, h, j), and the data is based on n=808 (a, d-f) n=408 (b-c), n=800 (g-j). Statistic based on unpaired T-test (a, c, h), Pearson r correlation and simple linear regression line (d), and calculated in Depmap (e, i).

Finally, we explored if specific gene expression patterns could predict if the CRISPR-p53 axis is active in a cell, and thus if the cell would enrich for TP53 sgRNAs. Based on our data identifying a central, non-redundant role for CDKN1A in the CRISPR-induced response, we tested if a strong baseline expression of *CDKN1A* could predict if cells would enrich for TP53 sgRNAs. We found that this assumption was correct and that cells that enrich for TP53 sgRNAs with a few exceptions had a strong baseline *CDKN1A* expression (**Fig. 3g-h**), while, for example, *TP53* expression itself did not predict cells that would enrich for TP53 sgRNAs (**Supplementary Fig. 9a-b**). Correlating strong baseline gene expression with TP53 sgRNA enrichment, resulted in a list of genes that to a large extent were identified as transcription target genes for p53 (**Fig. 3i**), and these genes were also identified by geneMANIA to display a strong co-expression pattern (**Supplementary Fig. 9c-f**). Notably, the gene set we identify by analyzing the 800 cell lines overlaps to a large extent with published gene expression patterns induced by CRISPR-mediated DNA damage in specific cell lines^18,46^, as well as genes upregulated in cancers WT for *TP53*^47^. Even more, performing a tSNE dimensional reduction analysis based on the expression of the top 10 genes, identified that cells broadly clustered based on *TP53* mutation status and TP53 sgRNA enrichment (**Fig. 3j** **and** **Supplementary Fig. 9g**). Interestingly, most of the cell lines that enriched for TP53 sgRNAs despite having *TP53* mutations also clustered with the *TP53* WT cells based on expression of the top 10 genes, suggesting that the specific mutations in those cell lines are not abrogating the p53 function sufficiently to cause a phenotype. Apart from *Cdkn1a*, the expression gene set list (Fig. 3i) does not overlap with genes identified to play a central role in the CRISPR-p53 pathway (Fig. 3e). This suggests that the upregulated genes are mainly an indication of active p53-mediated transcription in the cell, and not directly involved in the CRISPR response. *Eda2r* and *Ptchd4* knockout cells supported this notion by not behaving differently than WT cells in response to CRISPR in a mixed cell population (**Supplementary Fig. 9h-i)**. This observation challenges the notion that p53 is only active during stress, by showing that a baseline p53-regulated gene expression pattern before exposure to the specific stressor, CRISPR in this case, predicts the subsequent p53-related response to the stressor.

In conclusion, here we set out to explore if and how cells with inactive p53 are enriched as CRISPR is applied to a cell population, something that could represent a challenge for the clinical use of CRISPR. We identify that cells with mutations in *p53* indeed are enriched as CRISPR is applied, and that it correlates to the level of induced DNA damage response, highlighting the induction of *CDKN1A* expression as a sgRNA selection criterion. Furthermore, we identify a core CRISPR-p53 interactome, with genes that display a large level of physical interaction. Mutations or duplications of these genes could be enriched as CRISPR is applied, and these genes should, therefore, be monitored in the clinical CRISPR setting. Transient p53 inhibition has been proposed as a strategy to increase CRISPR efficiency^18–20^ and to retain the functionality of targeted cells^46,48^. Significantly, our data also show that transient p53 inhibition can limit the enrichment of mutations in the CRISPR-p53 tumor suppressor interactome, further supporting using transient p53 inhibition in the clinical CRISPR setting. Finally, we identify a gene expression profile, including strong baseline *CDKN1A* expression, that defines cells where the CRISPR-p53 axis is active.

## Methods

Methods, including statements of data availability and any associated accession codes and references, are available as supplemental methods.

## Acknowledgments

We are grateful to Drs. Robert M. Anthony, Kate Jeffrey, Eduardo Villablanca, and Bernhard Schmierer for their critical review of the manuscript, as well as to Drs. Michael Sundström, Lars Klareskog, Rasmus O. Bak, Alexander Espinosa, Klas G. Wiman, Lisa Westerberg, and Richard Rosenquist Brandell for important suggestions. psPAX2 (Addgene plasmid 12260) and pMD2.G (Addgene plasmid 12259) were kind gifts from Dr. Didier Trono. The ER-Hoxb8, and EcoPac plasmids were kind gifts from Dr. Mark P. Kamps. The R-script used to represent individual sgRNAs in Fig. S4a-b was a kind gift from Dr. Eric Shifrut. This work benefitted from data assembled by the DepMap Consortium for which we are grateful. The authors acknowledge support from the National Genomics Infrastructure in Stockholm funded by Science for Life Laboratory, the Knut and Alice Wallenberg Foundation and the Swedish Research Council, and SNIC/Uppsala Multidisciplinary Center for Advanced Computational Science for assistance with massively parallel sequencing and access to the UPPMAX computational infrastructure. This research was partly funded by grants from the Swedish Research Council (FW and DPL), the Swedish Cancer Society, Karolinska Institutet, Åke Olssons stiftelse (to FW), the China Scholarship Council (LJ and YS), and the Nanyang Technological University–Karolinska Institutet Joint PhD Programme (VSI).

## Author contributions

LJ and FW designed experiments with input from KI, SS and DPL. LJ and KI performed experiments. LJ, KI, YS, SVB, VSI, ZK, SS, DPL, and FW analyzed the data. LJ, DPL, and FW wrote the manuscript. The final manuscript was read and approved by all the authors.

## Competing interest statement

None declared.

## Online Methods

### Cells

The Hox bone marrow cell line was generated by transducing bone marrow cells of C57BL/6 Cas9+ GFP+ mice (The Jackson Laboratory, #026179) with an estrogen inducible retroviral construct expressing HoxB8 (ER-Hoxb8, a kind gift from Mark P. Kamps, University of California, San Diego) as described in^1,2^. Cells were cultured in 1 μM β-estradiol (BE, Sigma-Aldrich #E2758) and 25 nM mouse Stem Cell Factor (SCF, Peprotech #250-03) for several weeks with HoxB8 expression turned on to establish a cell line like population. Hox cells were cultured in RPMI-1640 (Sigma-Aldrich #R0883) with 10% heat-inactivated fetal bovine serum, 1% penicillin-streptomycin-glutamine (Gibco #10378016), 1 μM BE, and 25 nM SCF.

B16-F10 cell is a mouse melanoma cell line, purchased from ATCC (#CRL6475) and used at a low passage number. Cas9 expressing cells were generated by transducing B16-F10 cells with lentiCas9-Blast (Addgene #52962) lentiviral particles. B16 cells were cultured in RPMI-1640 (Sigma-Aldrich #R0883) with 10% heat-inactivated fetal bovine serum, and 1% penicillin-streptomycin-glutamine (Gibco #10378016)

### Viral preparation and transduction

Lentiviral particles were generated by seeding 2×10^6^ HEK293T cells in 10 cm plate in 10 ml of DMEM (Sigma-Aldrich #D6546) with 10% heat-inactivated fetal bovine serum and 1% L-glutamine (Gibco #A2916801). After ~24 h of culture, the cells reached ~60-70% confluency, and the medium was replaced by 5 ml of prewarmed fresh media. Transfer plasmids (lentiCas9-Blast, Addgene #52962; or LentiGuide-Puro-P2A-EGFP_mRFPstuf, Addgene #137730), pMD2.G (Addgene #12259), and psPAX2 (Addgene #12260) were mixed at 4:5:1 ratio (10 μg : 12.5 μg : 2.5 μg for 10 cm plate), and transfected into HEK293T cells using Lyovec (Invivogen #lyec) according to the manufacturer’s protocol. After 12 h, the medium was replaced by 8 ml of DMEM with 30% heat-inactivated fetal bovine serum and 1% L-glutamine. After another 36 h, the supernatant containing the virus was collected and centrifuged to remove the cell debris and used to spin infect cells.

For ER-Hoxb8 retrovirus preparation, plasmids including ER-Hoxb8 and the EcoPac gag-pol-env plasmid (both are kind gifts from Mark P. Kamps, University of California, San Diego) were mixed at 1:1 ratio (12 μg:12 μg for 10 cm plate) for transfection into HEK293T following the same approach as for generating lentiviral particles.

To transduce Hox or B16 cells with LentiGuide-Puro-P2A-EGFP_mRFPstuf, the multiplicity of infection (MOI) of the viral particles was tested by infection with serial dilutions of virus particles, to find a dilution resulting in a suitable MOI described below. Virus supernatant was added to each well of 6-well plate containing cells (4×10^5^ for Hox cells with SCF and BE, and 1×10^5^ for B16 cells) with 8 μg/ml polybrene (Sigma-Aldrich #H9268). The plate was centrifuged at 37°C, 1200g (120 min for Hox cells, and 45 min for B16 cells). After 24h, the virus-containing medium was replaced with fresh medium, and the infection rate was measured by the percentage of GFP+ cells if the vector contains GFP. Puromycin (Invivogen #ant-pr) selection (10 μg/ml for HoxB8 cells, and 5ug/ml for B16 cells) or Blasticidin (Invivogen #ant-bl) selection (10 μg/ml for B16 cells) was performed for 24 h to remove the non-infected cells.

### Electroporation and transfection

sgRNAs were designed using the Green Listed software^3,4^ using sgRNA design from the Doench mouse library^5^. 2’-O-methyl and phosphorothioate stabilized sgRNAs (**Supplementary Table 8**) were ordered from Sigma-Aldrich. Off-target activity was calculated using Cas-OFFinder (http://www.rgenome.net/cas-offinder/)^6^, and On-target activity was extracted from CHOPCHOP (https://chopchop.cbu.uib.no/)^7^ by searching for Ccr1, or entering the EGFP FASTA sequence from the LentiGuide-Puro-P2A-EGFP_mRFPstuf (Addgene #137730).

For Hox cells, Neon Transfection System (Invitrogen #MPK5000) was used to perform the electroporation following the manufacturer’s instructions (Pulse voltage: 1700, Pulse width: 20 ms, Pulse number: 1, for Hox cells). 100 pmol of sgRNA were electroporated into 2×10^5^ Hox cells for each electroporation experiment using Neon Transfection System 10 μL Kit (Invitrogen #MPK1096). For B16 cells, Lipofectamine 2000 Transfection Reagent (Invitrogen, #11668019) was used following the recommended protocol. 100pmol of sgRNA were transfected into 1×10^5^ B16 cells for each transfection experiment.

Trp53 ON-TARGETplus siRNA SMARTPool was ordered from Horizon. For Hox cells, 100 pmol of siRNA was electroporated into 2×10^5^ Hox cells for each electroporation experiment. For B16 cells, 100 pmol of siRNA was transfected into 1×10^5^ B16 cells for each transfection experiment. The Trp53 siRNA was typically delivered in the same reaction as the sgRNAs.

### CRISPR KO genotyping

1×10^5^ cells were collected for genomic DNA extraction using DNeasy Blood & Tissue Kit (Qiagen #69504) following the recommended protocol. Primers (Sigma-Aldrich) were designed using Primer-BLAST (**Supplementary Table 8**), aiming for a 400-1000 bp amplicon with the sgRNA target in the middle. Amplicons were gel purified and recovered using Zymoclean Gel DNA Recovery Kit (Zymo Research #D4007/D4008). The PCR products were quantified using Nanodrop and sequenced by Eurofins Genomics. The Sanger sequencing data was subsequently analyzed by ICE (Synthego, https://ice.synthego.com).

### Growth curve characterization

Hox cells were cultured with the following interventions: electroporation with a GFP targeting sgRNA; 0.5 μg/ml Etoposide (Sigma-Aldrich #E1383); 3 μM AMG232 (Axon Medchem #2639). The Etoposide and AMG232 were removed after 8 h by removing the medium and adding fresh medium without Etoposide or AMG232. The cells were cultured in 6 well plates with 4ml medium and passaged every day at the ratio of 1:3. 80 ul of cells were taken every day for flow cytometry (BD Accuri) with the existence of CountBright Absolute Counting Beads (Invitrogen #C36950). The absolute viable cell number was calculated by comparing the events number of viable cells and counting beads in the flow cytometry data.

### Real-Time PCR

TRIzol Reagent (Invitrogen #15596026) and Direct-zol RNA MiniPrep Kit (Zymo Research #R2051) was used to extract RNA. RNA was then converted into cDNA using High Capacity RNA-to cDNA kit (Applied Biosystem #4388950). The expression of Cdkn1a was quantified with a CFX 384 Real-Time PCR machine (Bio-Rad) using TaqMan gene expression FAM assays for Cdkn1a (Mm00432448_m1) with the TaqMan Gene Expression Master Mix (Applied Biosystem #4369542) as suggested by the manufacturer. Expression was normalized by TaqMan gene expression VIC assays for β-actin (Mm00607939) and Gene expression was quantified using the ddCT method.

### Apoptosis TUNEL assay

Cells were collected and fixed by PFA at different time point, and FlowTAC Apoptosis Detection Kit (R&D Systems #4817-60-K) was used to stain apoptotic cells following the manufacturer’s instructions. Stained cells were analyzed by flow cytometry (BD Accuri).

### Cloning of sgRNAs into lentiviral transfer plasmid, and CRISPR screens

sgRNAs with overhangs for the plasmids (**Supplementary Table 1**) were designed using the Green Listed software^3,4^ using sgRNA design from the Doench mouse library^5^ and, for intergenic controls, the Wang mouse library^8^. Individual sgRNAs were ordered from Sigma-Aldrich, and the sgRNA library was ordered from CustomArray as a DNA oligo pool. Cloning was performed using 150 ng of BsmBI (New England Biolabs #R0739) cleaved lentiGuide-Puro-P2A-EGFP_mRFPstuf plasmid (Addgene #137730), and 10 ng of the library oligo pool, using NEBuilder HiFi DNA assembly master mix (New England Biolabs #E2621). Endura ElectroCompetent Cells (Lucigen #60242), were subsequently transformed with the cloned plasmid pool using electroporation (1.0 mm cuvette, 10 μF, 600 Ohms, 1800 Volts) following the suggested protocol. The electroporated cells were combined and seeded on ten 20 cm LB agar plates with 100 μg/ml carbenicillin and grown at 37°C overnight. The lentiGuide-Puro-P2A-EGFP_mRFPstuf plasmid (Addgene #137730) makes bacteria red if the stuffer has not been exchanged by a sgRNA, and a few red clones were removed before collecting all other white clones. Then the plasmids were purified using EndoFree Plasmid Maxi Kit (Qiagen #12362).

The sgRNA cloned lentiGuide-Puro-P2A-EGFP_mRFPstuf was used as transfer plasmid for lentiviral preparation and transduction. The total amount of transduced cells was calculated based on MOI (0.25 for B16 cells, and 0.05-0.1 for Hox cells, which were difficult to transduce), based on the % GFP+ cells before puromycin selection, aiming for 1000 transduced cells for each sgRNA.

The CRISPR library transduced cells were exposed to GFP targeting sgRNA electroporation with or without Trp53 siRNA, 0.5 μg/ml Etoposide (Sigma-Aldrich #E1383) 8 h pulse stimulation, or 3 μM AMG232 (Axon Medchem #2639) 8 h pulse stimulation.

Cells were collected for genomic DNA extraction using DNeasy Blood & Tissue Kit (Qiagen #69504). Genomic DNA was then amplified using Q5 High-Fidelity DNA Polymerase (New England Biolabs #M0491) while introduced sample-specific barcodes and adapters for Illumina Sequencing similar as described in Joung J et al.^9^ using primers specified in **Supplementary Table 9**. The final PCR products were gel purified and recovered using Zymoclean Gel DNA Recovery Kit (Zymo Research #D4007/D4008), and quantified with Qubit 4 Fluorometer (Invitrogen #Q33238) using Qubit dsDNA HS Assay Kit (Invitrogen #Q32851) and pooled for next-generation sequencing (Illumina MiSeq v3 run, 2×75bp reads). The raw FASTQ data were analyzed by MAGeCK^10^. Read counts from CRISPR screens found in **Supplementary Table 10-11**.

### JAK1/STAT1 signaling assay

Hox cells were cultured with or without mouse Interferon Beta (IFNβ, Nordic BioSite #12405-1) and the Jak1 inhibitor Solcitinib (Selleckchem #S5917) for 7 days. Cells were passaged at the ratio of 1:8 every other day. 100 ul of cells were taken on day 7 for flow cytometry (BD Accuri) with the existence of CountBright Absolute Counting Beads (Invitrogen #C36950). The absolute viable cell number was calculated by comparing the events number of viable cells and counting beads in the flow cytometry data.

### Competitive co-culture assay

Trp53 KO and WT cells were mixed at 1:4 ratio, and were electroporated with sgRNA or transduced with lentivirus (lentiGuide-Puro-P2A-EGFP_mRFPstuf) and cultured for 7 days; or cultured with Etoposide (Sigma-Aldrich, #E1383, 0.05 μg/ml) or AMG232 (Axon Medchem #2639, 0.5 μM for Hox cells, 4 μM for B16 cells) for 7 days; or cultured with Cobalt(II) chloride (CoCl_2_, Sigma-Aldrich #232696, 10-20 μg/ml) for 7 days. Different p53 related inhibitors were added during culture: Trp53 ON-TARGETplus siRNA SMARTPool (Horizon), KU55933 (Sigma-Aldrich #SML1109, 100 ng/ml for B16, 10 ng/ml for Hox), VE821 (Sigma-Aldrich #SML1415, 50 ng/ml), Pifithrin-μ (Sigma-Aldrich #P0122, 2 μg/ml for B16, 200 ng/ml for Hox), Cyclic Pifithrin-α (Sigma-Aldrich #P4236, 700 ng/ml for B16, 70 ng/ml for Hox), C646 (Sigma-Aldrich, #SML0002, 1 μg/ml), AZD2461 (Selleckchem #S7029, 250 ng/ml), LJI308 (Sigma-Aldrich #SML1788, 2 ng/ml), Z-VAD-FMK (Selleckchem #S7023, 50 μM). siRNA was delivered to cells 1 day before CRISPR/Etoposide/AMG232 exposure (100 pmol of siRNA electroporated into 2×10^5^ Hox cells, or transfected into 1×10^5^ B16 cells), or together with sgRNA for the transfection/electroporation groups. Other inhibitors were added to cell culture media 1 day before CRISPR/Etoposide/AMG232 exposure and cultured for 7 days. Cells were then collected for Trp53 KO genotyping. **Supplementary Table 7** further describes how stock solutions of inhibitors were generated and stored.

### Flow Cytometry Analysis

Fresh bone marrow cells from C57BL/6 Cas9+ GFP+ mice and Hox cells were stained with the following antibodies: FITC Rat anti-Mouse CD34 (BD Biosciences #553733, 1:500), PE anti-mouse CD150 (BioLegend #115904, 1:200), PerCP/Cyanine5.5 anti-mouse Ly-6A/E (BioLegend #122524, 1:200), APC anti-mouse CD117 (BioLegend #105812, 1:500), APC/Cyanine7 anti-mouse CD16/32 (BioLegend #156612, 1:500), Biotin anti-mouse Lineage Panel (BioLegend #133307, 1:100), BV421 Streptavidin (BD Biosciences #563259, 1:1000), LIVE/DEAD Fixable Aqua Dead Cell Stain Kit (Invitrogen #L34957, 1:2000). After 30 min of staining, the cells were washed and analyzed by flow cytometry (BD FACSVerse). FACS FCS files were analyzed by FlowJo version 10 (FlowJo, LLC).

### Analysis of data from the Depmap portal

sgRNA enrichment (CRISPR (Avana) Public 20Q4 release), mutation profile (Mutation Public 20Q4 release), drug sensitivity (PRISM Repurposing Primary Screen 19Q4 release), and mRNA expression levels (Expression Public 20Q4 release) was extracted December 13^th^ 2020 from the Depmap portal (https://depmap.org/portal/)^11–13^. Connectivity maps were generated using the geneMANIA plugin for Cytoscape^14,15^. tSNE plots were made with the Rtsne package (https://github.com/jkrijthe/Rtsne) to analyze the cluster and ggplot2 (https://github.com/tidyverse/ggplot2) to visualize the data, or tSNE-online (https://github.com/jefworks/tsne-online). The “ENCODE and ChEA Consensus TFs from ChIP-X” functionality of Enrichr (https://maayanlab.cloud/Enrichr/index.html)^16,17^ was used to identify transcription factor binding to gene sets.

---

**Supplementary Figure 1.**
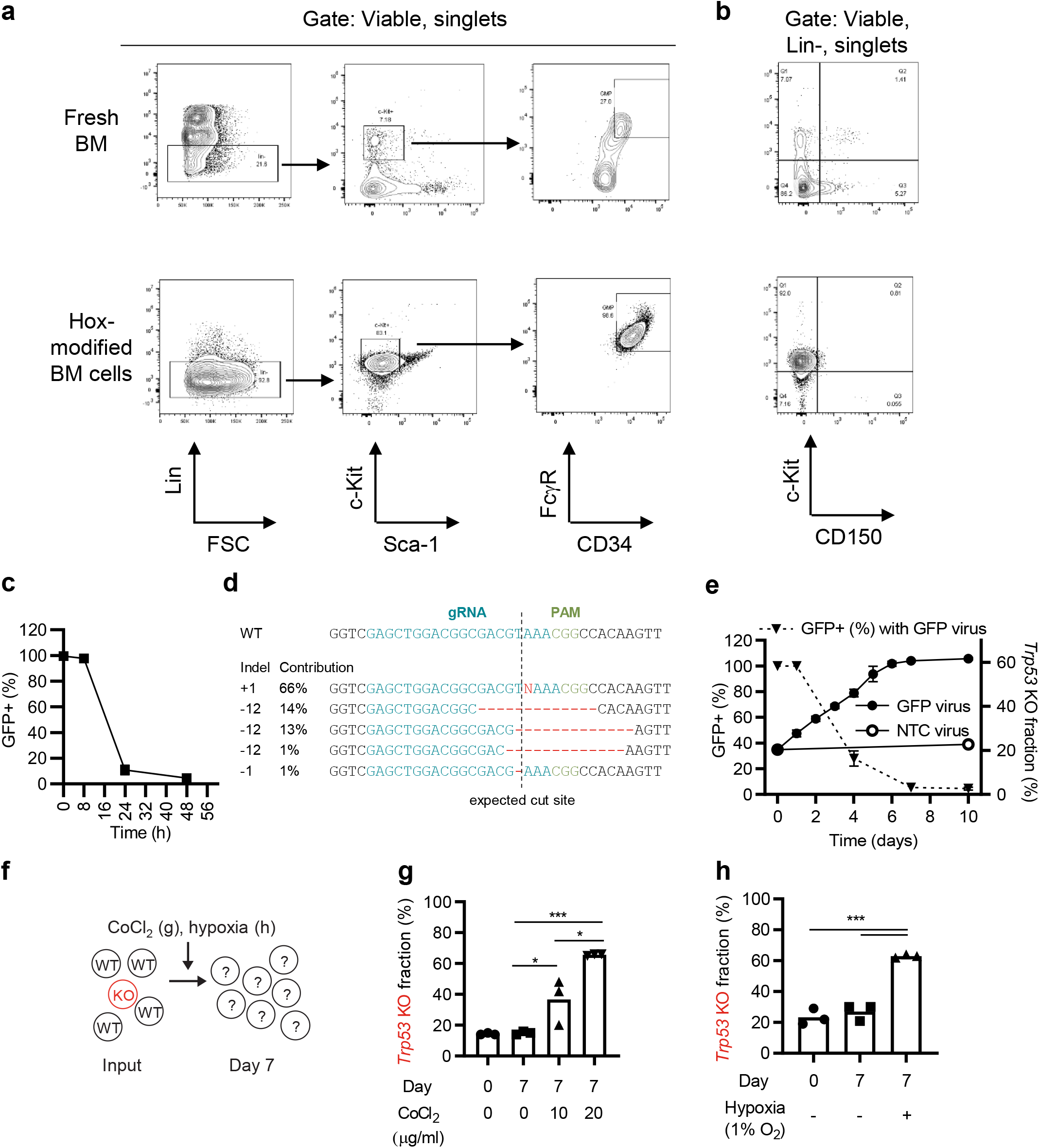
Hox cells, CRISPR-mediated GFP inactivation, and hypoxia. (**a-b**) Flow cytometry analysis of fresh bone marrow (BM) and Hox expanded BM cells from C57BL/6 Cas9+ GFP+ mice. Hox cells display a Granulocyte-Monocyte Progenitor (GMP) phenotype (Lin-, c-kit+, Sca-1-, FcgR+, CD34+, CD150-). (**c**) Kinetic flow cytometer analysis of GFP signal of Hox cells electroporated with a GFP targeting sgRNA. (**d**) Sanger sequencing and ICE mutation analysis of the GFP targeted region in Hox cells 48h after electroporation of a GFP targeting sgRNA. (**e**) Kinetic flow cytometer analysis of GFP signal of Hox cells transduced with lentiviral particles carrying a GFP targeting sgRNA. Triangles indicate the kinetics of GFP KO (left y-axis), showing a substantially slower dynamics compared to sgRNA electroporation in (c). Circles indicate the kinetics of the enrichment of Trp53 KO cells (right y-axis). (**f-h**) WT and Trp53 KO Hox cells were mixed and cultured for seven days in the presence of different concentrations of Cobalt Cloride (CoCL_2_) that stabilizes hypoxia-inducible factor-1α (g) or in a hypoxia chamber (h). Data presented as mean +/− SEM, n=3 (c, e), or mean and individual values (g-h). Data is representative of two or more experiments.

**Supplementary Figure 2.**
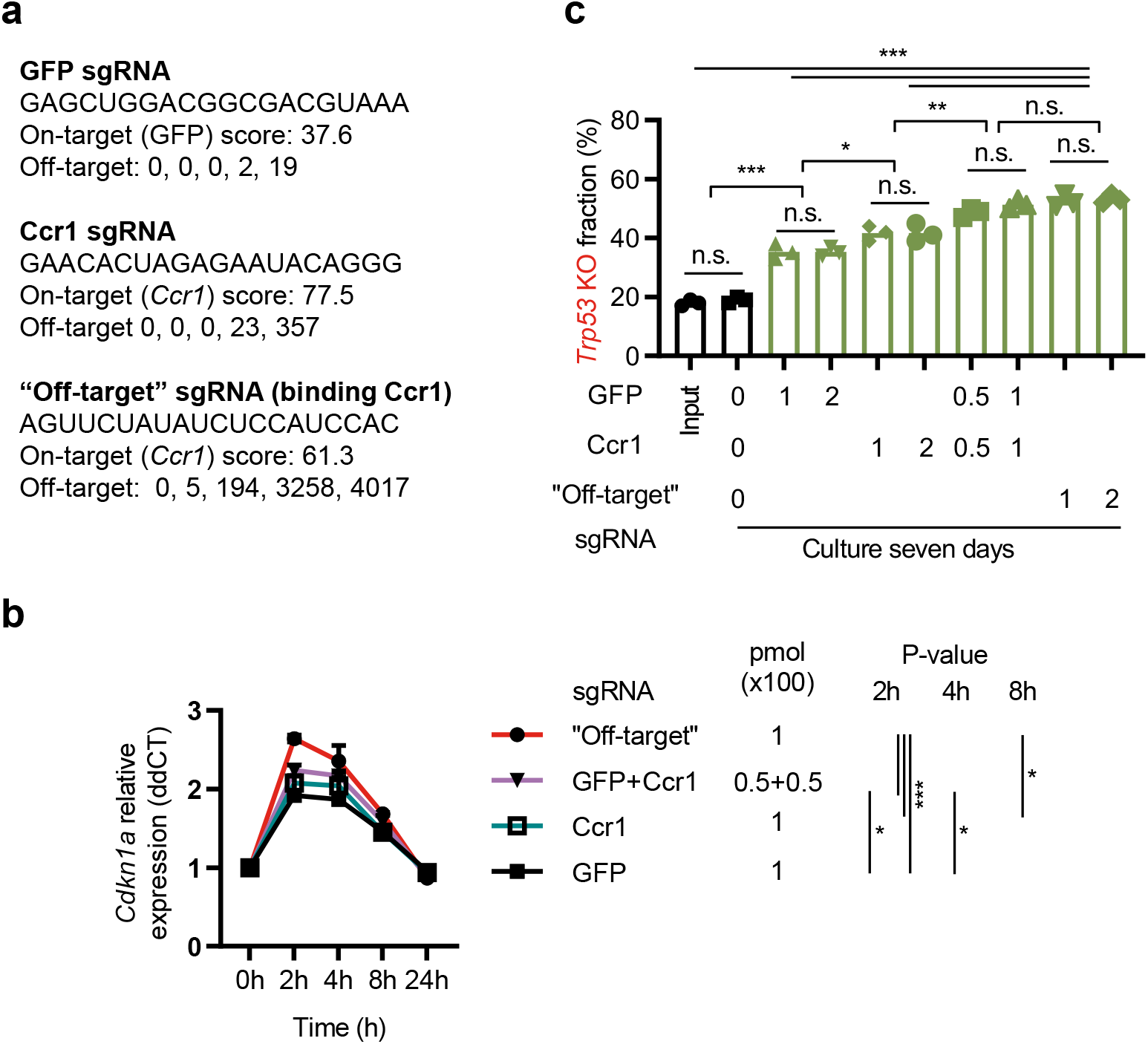
The level of DNA damage induced by CRISPR affects the magnitude of enrichment of cells with *Trp53* mutations. (**a**) Information about sgRNAs used in the figure. On-target score calculated by CHOPCHOP (PMID: 31106371). Off-target indicates number of targets with 0, 1, 2, 3, and 4 miss matches to the mouse genome as identified by Cas-OFFinder. (**b**) Hox cells (Cas9+ and GFP+) were electroporated by different sgRNAs at the indicated doses (pmol x 100), and cells collected for *Cdkn1a* qPCR (same data as shown in heatmap, Fig. 1g, but with quantification). (**c**) *Trp53* KO and WT Hox cells (Cas9+ and GFP+) were mixed 1:5 and electroporated by different sgRNAs at the indicated doses (pmol x 100). Seven days later cells were collected and sequenced to determine the % *Trp53* KO sequences. Data presented as mean +/− SEM, n=3 (b), or mean and individual data, n=3 (c). Data is representative of two or more experiments. * = p < 0.05, ** = p < 0.01, *** = p < 0.001, and n.s. = non-significant by two-way ANOVA and Tukey’s post-test (b), or one-way ANOVA and Tukey’s post-test (b).

**Supplementary Figure 3.**
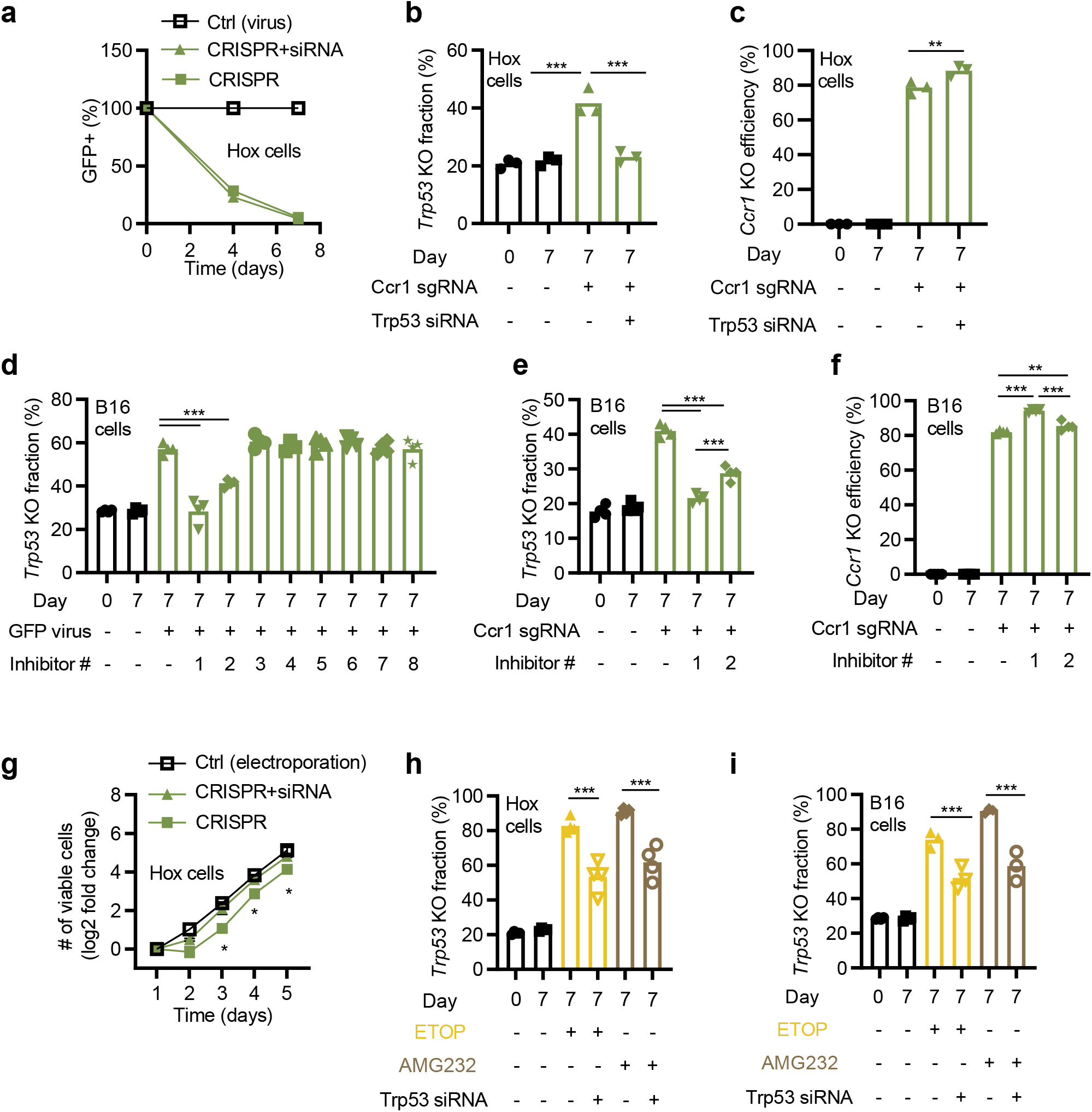
Transient inhibition of p53 limits enrichment of cells with mutations in *Trp53* while retaining, or increasing, the KO efficiency. (**a**) Hox cells (Cas9+ and GFP+) were electroporated +/− Trp53 siRNA and subsequently transduced with a virus carrying a GFP targeting sgRNA. The KO efficiency of GFP was followed over time by flow cytometry. (**b**) WT and *Trp53* KO Hox cells were mixed and electroporated with a Ccr1 sgRNA +/− Trp53 siRNA. Cells were cultured for seven days, followed by sequencing of Trp53 to quantify the frequency of mutations. (**c**) The *Ccr1* knockout efficiency was measured by sequencing *Ccr1* in Hox cells treated as in (b). (**d**) WT and p53 KO B16 cells (Cas9+ and GFP+), were mixed and transduced with a GFP targeting sgRNA virus in the presence of a selection of inhibitors (see figure 1i-j for details). Inhibitor #1 = Trp53 siRNA, #2 = KU55933 (an Atm inhibitor). Cells were then cultured for seven days, followed by sequencing of *Trp53*, and the frequency of *Trp53* mutations quantified. (**e-f**) Same experiment as (b-c), but with B16 cells. (**g**) Growth characteristics of Cas9+ GFP+ Hox cells electroporated with a GFP sgRNA +/− Trp53 siRNA. (**h-i**) WT and *Trp53* KO Hox cells (h) or B16 cells (i) were mixed and incubated with an 8h pulse of ETOP or AMG232 in the presence or absence of Trp53 siRNA. Data is shown as mean +/− SEM, n=3 (a, g), or mean and individual values, n=3-4 (b-c, d-f, h-i). Data is representative of two or more experiments. ** = P < 0.01, *** = P < 0.001 by one-way ANOVA and Tukey’s post-test (b-c, d-f, h-i), or two-way ANOVA and Tukey’s post-test (g, indicated significance relates to CRISPR +/− siRNA).

**Supplementary Figure 4.**
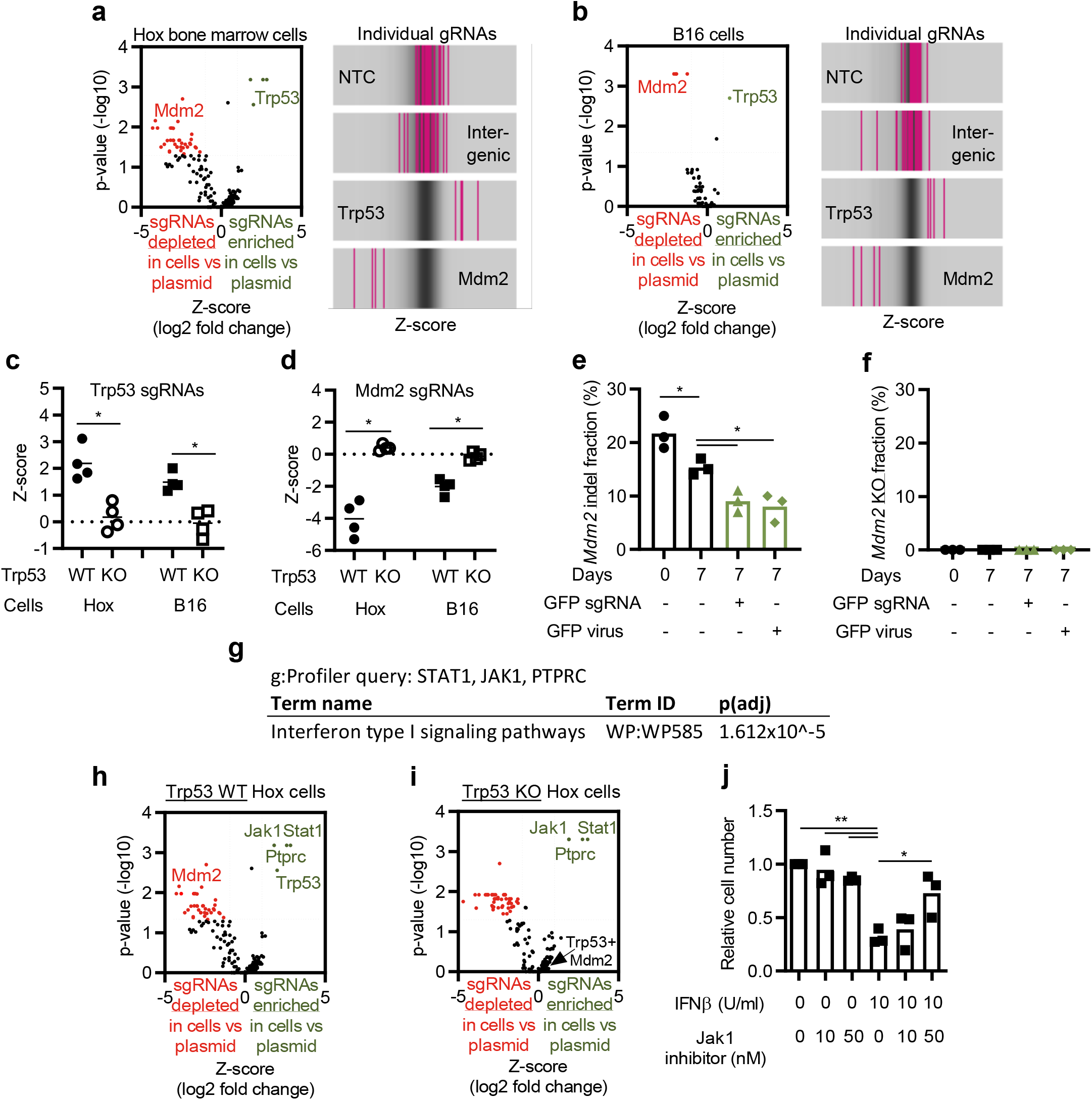
p53 and MDM2 play a central role in the CRISPR-mediated DNA damage response, and JAK1/STAT1 signaling negatively affects Hox cell survival independently of p53 status. (**a-b**) Exploratory CRISPR screen targeting 400 cell death-related genes and controls in Hox cells (a), and B16 cells (b). The sgRNA representation was analyzed by next-generation sequencing, and enrichment/depletion deconvoluted by MAGeCK. Genes (left) and individual sgRNAs of non-targeting controls (NTC), intergenic controls, Trp53, and Mdm2 (right) enriched and depleted after seven days by the CRISPR-induced DNA damage. (**c-d**) Enrichment and depletion of individual sgRNAs for Trp53 (c), and Mdm2 (d) in the CRISPR screen, comparing WT and Trp53 KO Hox and B16 cells. (**e-f**) Hox cells WT or with mutations (any Indels, including insertions and deletion of 3 nucleotides) in Mdm2 were mixed and exposed to CRISPR (sgRNA electroporation or sgRNA virus). Cells were cultured for seven days, followed by sequencing of Mdm2 to quantify the frequency of indels (e) and KO (f). (**g**) Top WP term identified by g:Profiler querying STAT1, JAK1, and PTPRC. (**h-i**) Enrichment of Jak1, Stat1, and Ptprc sgRNAs in Hox WT (h), and Hox Trp53 KO cells (i). (h) is the same data as (a), but indicating Jak1, Stat1, Ptprc. (**j**) Hox cells cultured for five days with type I interferon (IFNb) and Jak1 inhibitor. Data presented as relative cell number compared to the control (no IFNb or Jak1 inhibitor). Data is presented as volcano plots with adjusted p-values (−log10; >1.3 = p < 0.05) and Z-score (log2 fold enrichment/depletion of sgRNAs (a-b, g-h, adjusted p-value <0.05, and Z-score >1 or <−1 are indicated in color), or mean and individual values (c-f, j). * = p < 0.05, and ** = p < 0.01 by Mann-Whitney test (c, d), or one-way ANOVA and Turkey’s post-test (e-f, j).

**Supplementary Figure 5.**
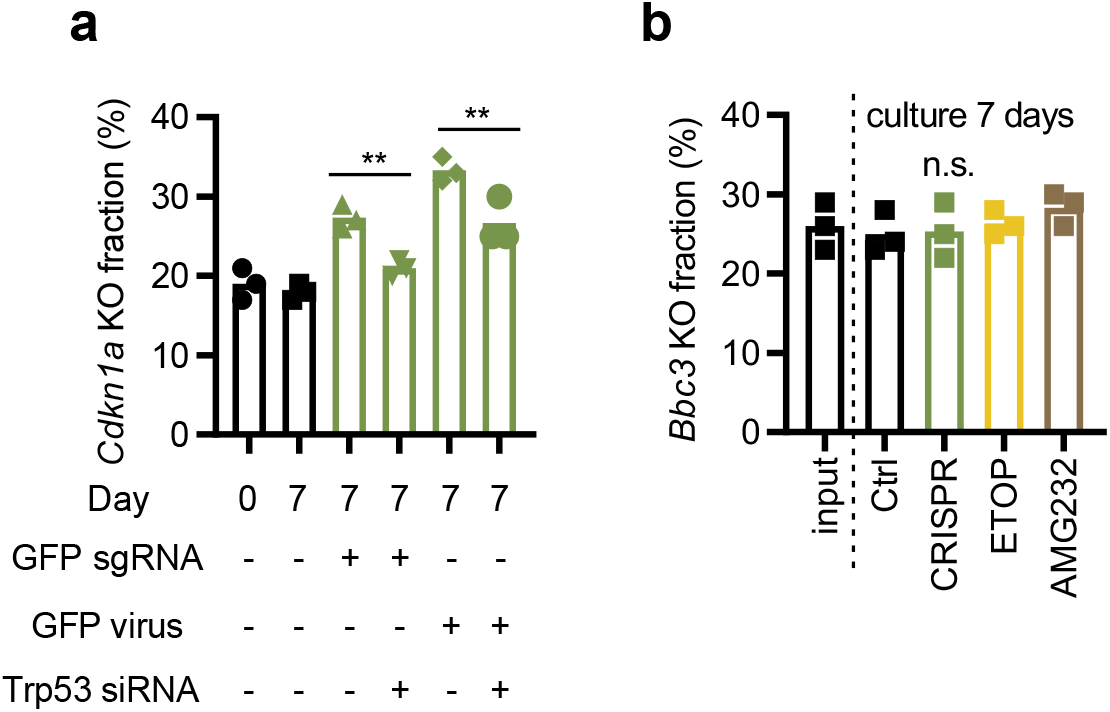
Enrichment of mutations in *Cdkn1a* but not *Bbc3* by CRISPR. (**a**) WT and *Cdkn1a* KO Hox cells were mixed and electroporated with a GFP sgRNA +/− Trp53 siRNA. Cells were cultured for seven days, followed by sequencing of *Cdkn1a* to quantify the frequency of mutations. (**b**) WT and *Bbc3* KO Hox cells were mixed and electroporated with a GFP sgRNA, or exposed to a 8h pulse with Etoposide or AMG232. Cells were subsequently cultured for seven days, followed by sequencing of *Bbc3* to quantify the frequency of mutations. Data is presented as mean and individual values (pooled from three independent experiments) n.s. = non-significant, and ** = p < 0.01 by one-way ANOVA and Turkey’s post-test.

**Supplementary Figure 6.**
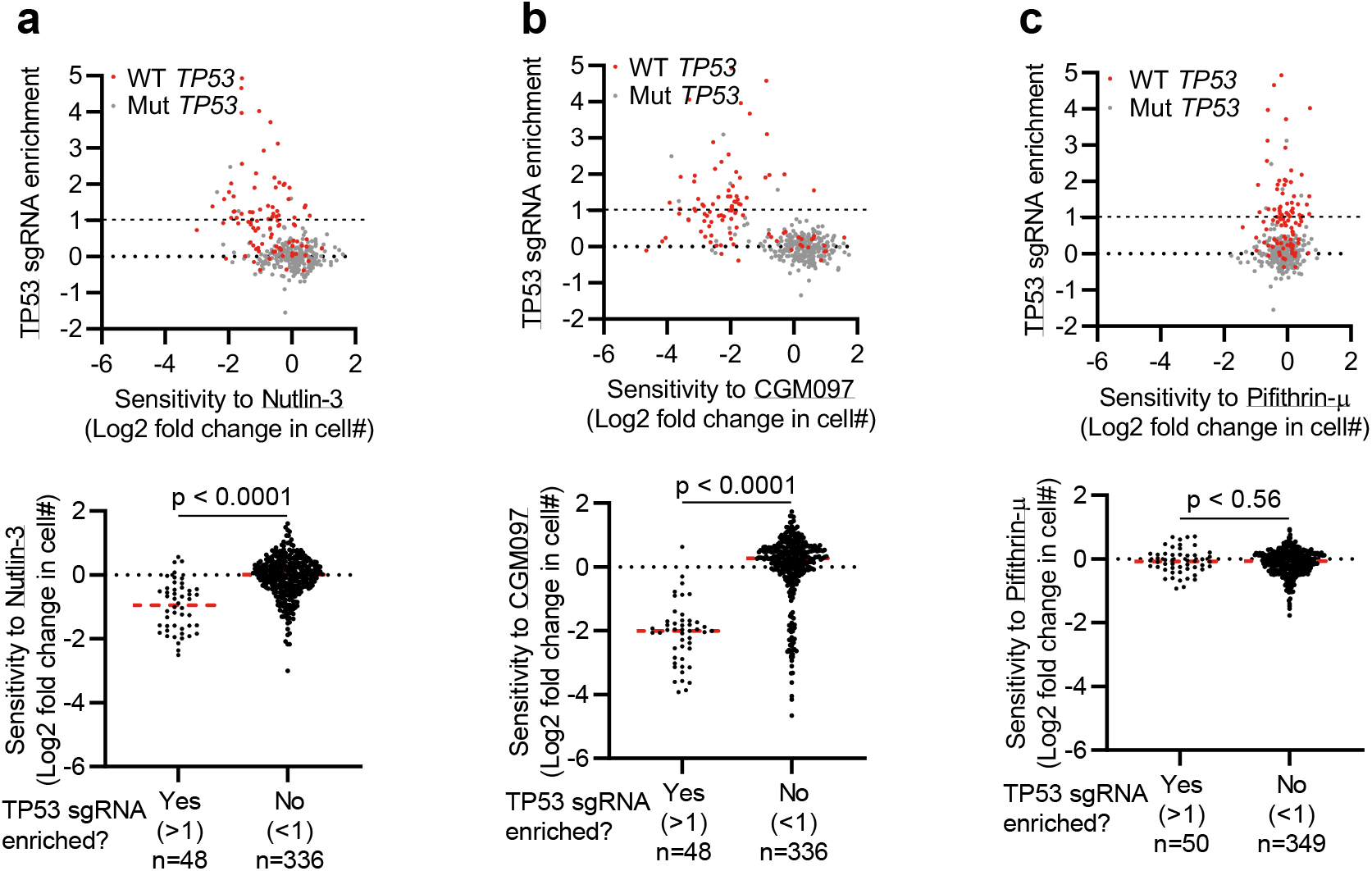
The sensitivity of cell lines to Nutlin-3, CMG097, and Pifithrin-m. (**a-c**) TP53 sgRNA enrichment score from the Depmap, and sensitivity to Nutlin-3 (a), CGM097 (b), and Pifithrin-m (c). Sensitivity is defined as the number of cells after a five-day culture, compared to control-treated cells (log2 fold change). Upper graphs show both parameters and *TP53* mutation status (WT or mutated, Mut) indicated by color, lower graphs show sensitivity in cells stratified based on TP53 sgRNA enrichment. Data includes all available data in the Depmap CRISPR (Avana) 20Q4, Mutation Public 20Q4, and Drug sensitivity (PRISM Repurposing Primary Screen) 19Q4 releases. Each dot represents one cell line. Statistics based on unpaired T-test (a-c).

**Supplementary Figure 7.**
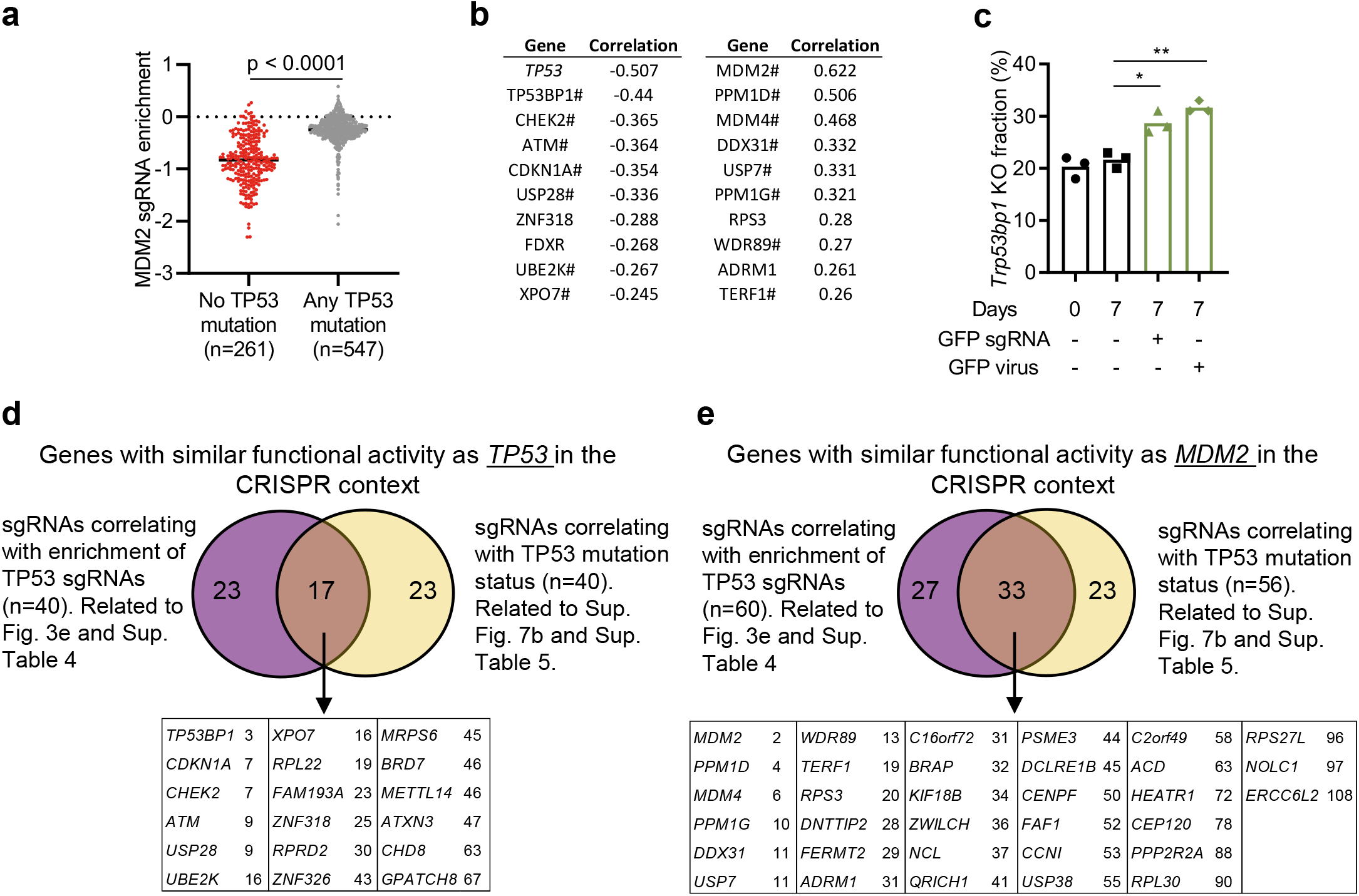
sgRNA enrichment/depletion that correlates with TP53 mutation status. (**a**) MDM2 gRNA enrichment score stratified based on *TP53* mutation status. (**b**) Top ten negative and positive correlating genes comparing sgRNA enrichment in cells that are WT for *TP53* or that has any mutation in *TP53*. # indicates genes that overlap with the sgRNA correlation list in figure 3e. (**c**) WT and *Trp53bp1* KO Hox cells were mixed and electroporated with a GFP sgRNA or transduced with a virus carrying the same GFP sgRNA. Cells were cultured for seven days, followed by sequencing of *Trp53bp1* to quantify the frequency of mutations. (**d-e**) Venn diagram showing overlap of genes identified to correlate with the two different approaches, divided into genes that functionally behave similarly as *TP53* (d), and *MDM2* (e), in the CRISPR context. Overlapping genes are sorted based on the combined ranking of the two different approaches. Data includes all available data in the Depmap CRISPR Avana 20Q4, and Mutation Public 20Q4 releases. Each dot represents one cell line. Statistics based on unpaired T test (a), correlation calculated by Depmap (b), or one-way ANOVA and Tukey’s post-test, * = p < 0.05, ** = p < 0.01 (c).

**Supplementary Figure 8.**
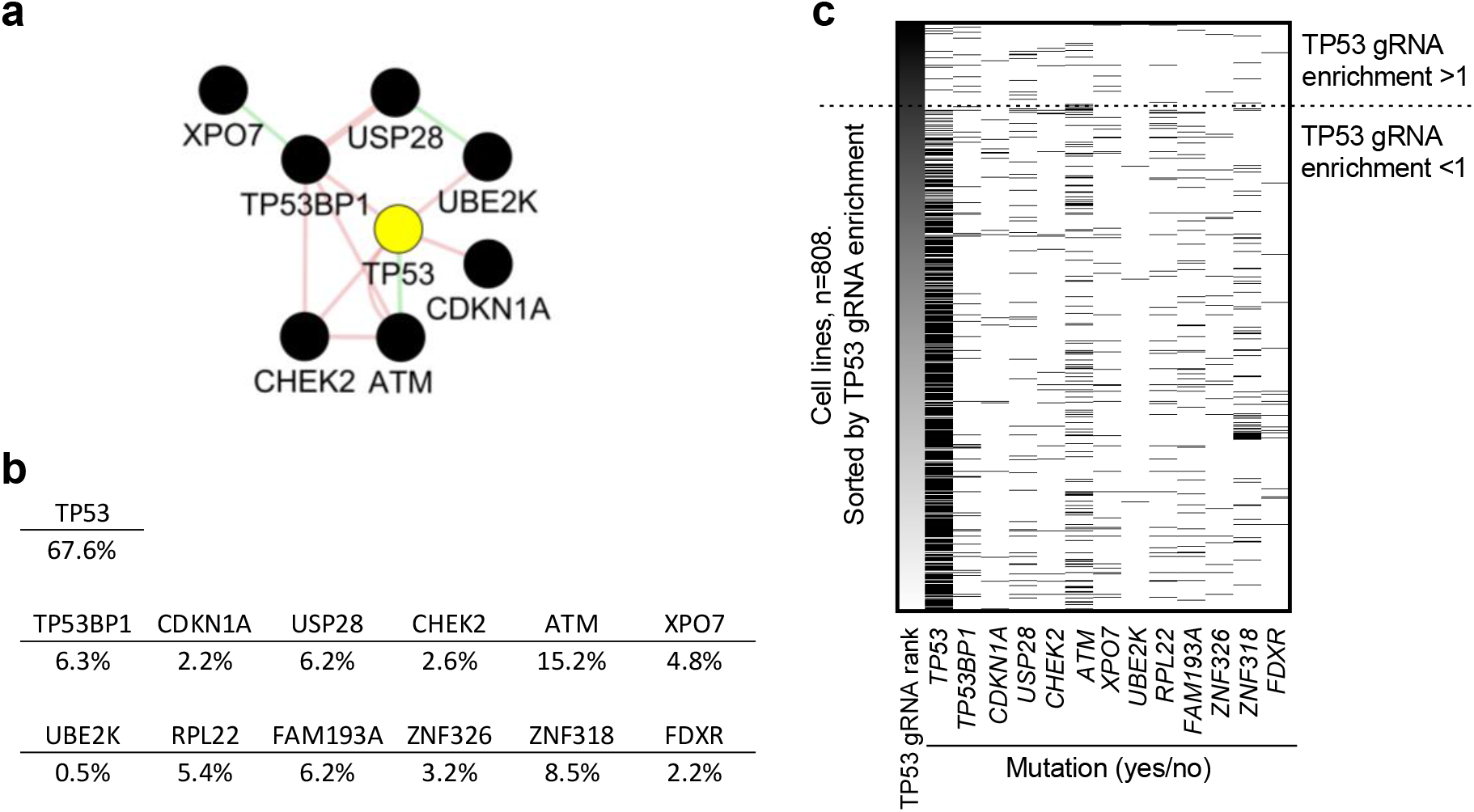
Mutation frequency in the core CRISPR-p53 tumor suppressor interactome. (**a**) A core CRISPR-p53 tumor suppressor interactome based on experimental data and analysis of Depmap correlations related to TP53 gRNA enrichment and *TP53* mutation. Includes tumor suppressor genes overlapping in Fig. 3e and Supplementary Fig. 6b. Interactions identified by geneMANIA: Red = physical interaction, green = genetic interaction. (**b**) Mutation frequency (% any mutation) in all cell lines where both CRISPR screen data and mutation data were available, n=808. Includes all genes identified in Fig. 3e and Supplementary Fig. 6b, the last five genes are, thus, only found in one of the top 10 lists. (**c**) Heat map showing mutations (white = WT, black = any mutation) of the indicated genes (all genes identified in Fig. 3E and Supplementary Fig. 6B). Data is based on all available data in the Depmap CRISPR (Avana) 20Q4, and Mutation Public 20Q4 releases.

**Supplementary Figure 9.**
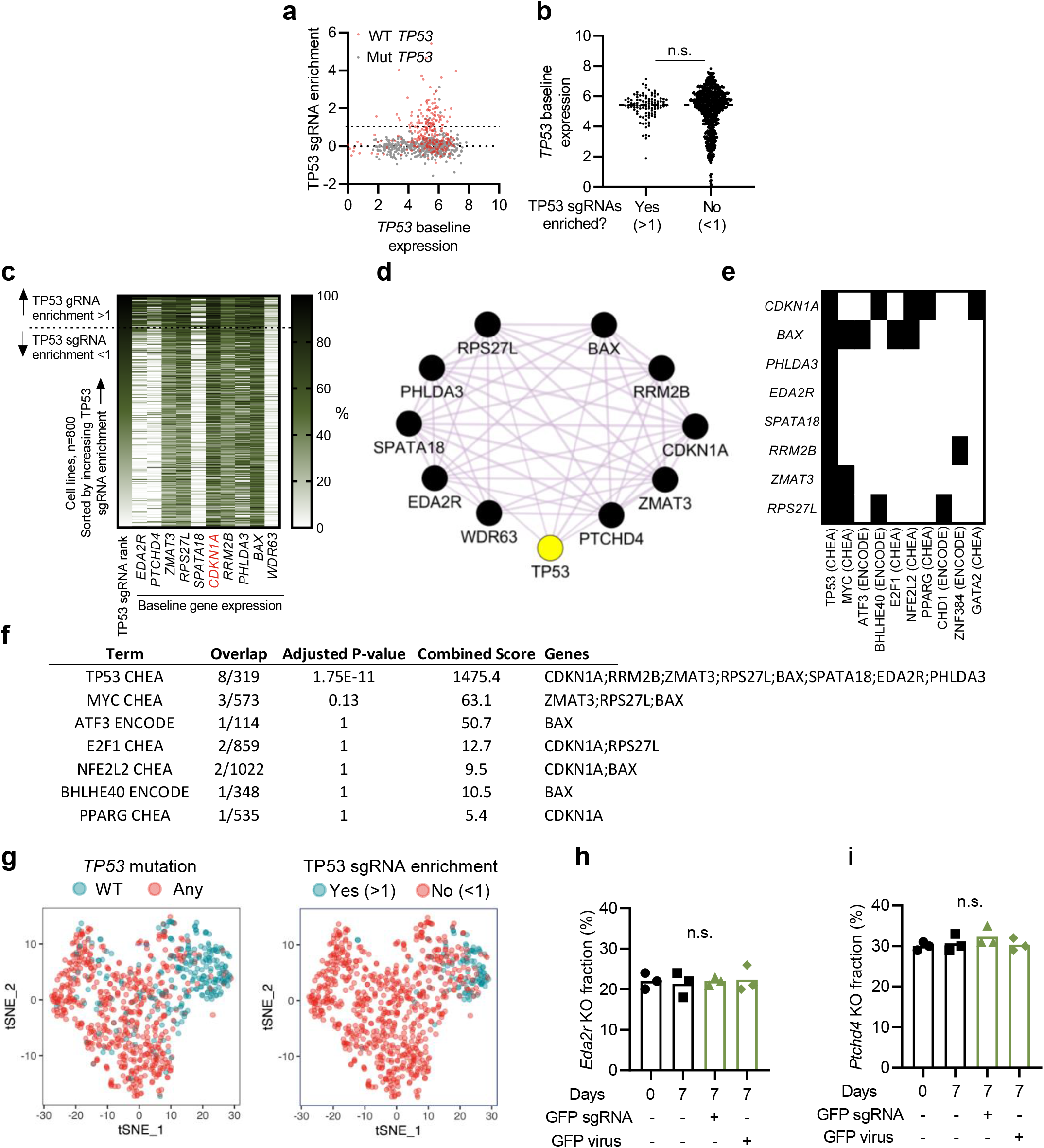
p53 target gene expression, but not *TP53* expression, predicts TP53 sgRNA enrichment. (**a-b**) Correlation between TP53 sgRNA enrichment and *TP53* baseline expression. (**c**) Baseline expression of the ten selected genes in 800 cell lines sorted by TP53 sgRNA enrichment rank. For visualization purposes, the expression is normalized to 0-100%, where the highest expression of each gene is defined as 100% and the lowest as 0%. (**d**) Interactions of the ten genes identified by geneMANIA: purple = co-expression. (**e-f**) Transcription factor binding to the top ten genes analyzed by the Enrichr software: ENCODE and ChEA Consensus TFs from ChIP-X. (e) Target genes (y-axis) for different transcription factors (TF, x-axis) indicated with black color. (f) Statistical analysis of as calculated by the Enrichr software of TFs linked to the gene set. (**g**) tSNE dimensionality reduction analysis of cells based on the expression of the 10 selected genes (same data as Fig. 3j but divided into two graphs for clarity). The colors indicate *TP53* mutation state (left) and TP53 sgRNA enrichment (right). (**h-i**) WT and *Eda2r* (h), or *Ptchd4* (i) KO Hox cells (Cas9+ and GFP+) were mixed and electroporated with a GFP sgRNA or transduced with a GFP sgRNA virus. Cells were cultured for seven days, followed by sequencing of *Eda2r* (h), or *Ptchd4* (i) to quantify the frequency of mutations. Data presented as individual cell lines (each dot/line represents one human cell line), n=800 (a-c, g), or as mean and individual data (h-i). Data includes all available data in the Depmap CRISPR (Avana), and Expression Public 20Q4 releases. Statistics based on unpaired T-test (b, h-i), n.s. = non-significant.

## Supplementary tables

1. sgRNA library.
2. Genes enriched/depleted in primary screen (related to Sup. Fig. 4a-d, g-i).
3. Cell lines included in Depmap with TP53 sgRNA enrichment (related to Fig. 3a).
4. Extended list of sgRNAs correlating to TP53 sgRNA enrichment (related to Fig. 3e).
5. Extended list of sgRNAs correlating to TP53 mutation status (related to Sup. Fig. 7b).
6. Extended list of gene expression correlating to TP53 sgRNA enrichment (related to Fig. 3i).
7. Reagents (stock solution and storage)
8. sgRNAs and primers used for electroporation.
9. Primers used for NGS analysis of CRISPR screens.
10. Read count first screen
11. Read count second screen

## Notes

### Competing Interest Statement

The authors have declared no competing interest.

